# Chaotic Dynamics and Effects of Cognitive State in Human Single Neurons

**DOI:** 10.1101/2025.08.10.669543

**Authors:** Saarth Anjali Chitale, Swetha Ravichandran

**Author notes:** **Materials & Correspondence** Correspondence and requests for materials should be addressed to S.A. Chitale.

## Abstract

How the brain balances flexibility and stability during memory-guided behavior remains unclear. While large-scale brain activity exhibits signatures of chaos, it is unknown whether individual neurons also operate in a chaotic regime, and whether such dynamics are modulated by task engagement. Here, we analyzed over 1800 medial temporal lobe (MTL) neurons recorded intracranially during a declarative memory task. Using sample entropy, correlation dimension, and the largest Lyapunov exponent, we found robust signatures of deterministic chaos in a substantial proportion of individual neurons. Crucially, these signatures were not static: during memory task engagement, neurons exhibited greater entropy and dimensionality, and lower divergence, compared to task-irrelevant periods. Chaos was widespread across MTL subregions and independent of age or gender. These results suggest that chaos is an intrinsic, dynamic feature of single-neuron activity which is selectively amplified during cognitive engagement, and may support flexible temporal coding in the human memory system.

## Introduction

How does the brain maintain both flexibility and stability during memory-guided behavior? One possibility is that neural systems operate in a dynamical regime known as low-dimensional chaos which is structured yet unpredictable, sensitive to input yet bounded in evolution. This hypothesis has been proposed to explain rapid transitions between cognitive states and the brain’s capacity for generalization, novelty detection, and internal exploration [1, 2, 3]. Although chaotic dynamics have been observed in macroscopic signals like EEG (electroencephalogram) and LFPs (local field potentials) [4, 5, 6, 7], it remains unknown whether chaos also emerges at the level of individual neurons in humans.

Theoretical models suggest that single neurons are capable of chaotic dynamics due to intrinsic nonlinearities in ion channel kinetics, synaptic integration, and bursting behavior [8, 9, 10]. Such dynamics could allow neurons to traverse representational spaces flexibly, supporting memory formation and recall [3, 11]. However, direct evidence of chaos in human single-neuron spike trains is scarce. Critically, even if chaos is present, it is unclear whether it reflects a passive biophysical feature or a dynamically regulated computational regime. Here, we hypothesize that chaotic structure in single-neuron spike timing is not static but is amplified during active cognitive engagement, particularly during memory tasks that require flexibility and internal trajectory evolution.

To investigate these possibilities, we applied a hybrid framework combining embedding-based and embedding-independent chaos metrics to a publicly available dataset of neuronal spike trains recorded from the medial temporal lobe (MTL) of epilepsy patients performing a declarative memory task [12]. We computed sample entropy (SE), correlation dimension (CD), and the largest Lyapunov exponent (LE) for interspike interval (ISI) sequences derived from the spike times. While SE was computed directly in ISI space making it embedding independent, both CD and LE were estimated via state-space embedding. To isolate genuine nonlinear structure or deterministic chaos from stochastic variability, we employed phase-randomized surrogate testing, ensuring conservative statistical inference by correcting for the maximum values across surrogate-derived distributions over parameter sweeps.

Across hundreds of neurons, we found robust and reproducible signatures of chaotic dynamics in a substantial subset. Chaos was more prevalent during active engagement in the task than during idle periods, suggesting functional modulation rather than random variability. The chaotic dynamics were spatially widespread across the MTL and not confined to any specific subregion. Moreover, presence of chaos did not vary significantly with age or gender. One exception is the statistically significant difference observed between genders during task-irrelevant period wherein greater proportion of neurons were chaotic in males compared to females. These re-sults suggest presence of deterministic chaos in human single-neuron activity during naturalistic behavior.

## Methods

The publicly available dataset [12] analyzed in this article was obtained from a study at three locations - Huntington Memorial, Cedars-Sinai, and Toronto Western Hospitals, with corresponding approvals from respective Institutional Review Boards. The data was recorded *in vivo* from 59 patients who were evaluated for potential surgical treatment of epilepsy as they performed a declarative memory task across 87 sessions. These recordings included the activities of single neurons in the MTL using an epilepsy monitoring unit. All participants provided written informed consent prior to the recordings and completed between one and three sessions, with new task variants and stimuli. This data is available in Neurodata Without Borders (NWB) format and can be accessed using the PyNWB interface. Spike times for individual neurons were obtained from the units table and mapped to trial boundaries using annotations in intervals/trials. Below, we give a brief description of the task along with the procedures for ISI extraction, chaos metric estimation, surrogate testing, and validation.

### Task and Data

The declarative memory task used in patients with pharmacoresistant epilepsy involved *Learn* and *Recognition* phases [12] as illustrated in Figure 1. During the *Learn* phase, subjects viewed images from various semantic categories and performed a binary animal detection task, indicating whether each image contained a `small animal`. In the *Recognition* phase, they were shown a mix of previously seen (in the *Learn* phase) or novel images and were asked to report whether they found the image to be familiar or new. The task design ensures that the *Learn* phase is visually-selective (VS) while the *Recognition* phase is memory-selective (MS), both of which are task-relevant phases. Some neural spikes in this dataset lie outside either *VS* or *MS* phases, and we presume these occur during the task-irrelevant (TI) period when participants were not actively engaged in the task.

**Figure 1:**
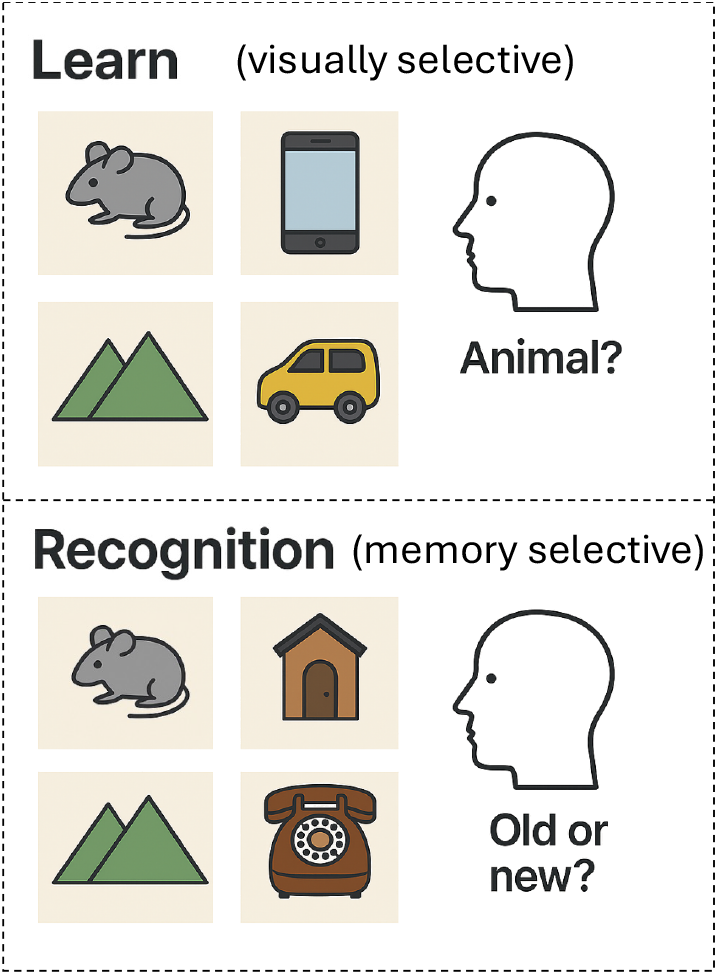
Visually- and memory-selective tasks used for neural recordings in epilepsy patients [12]. During visually-selective (*Learn*) phase, participants indicated whether each image contained a small animal. In contrast, during memory-selective (*Recognition*) phase, they judged whether each image had been presented previously (old) or was novel presentation (new).

The experimental setup described above yielded spike timing data for more than 1800 single neurons and to determine whether the activity of these neurons exhibits evidence of chaotic dynamics, we relied on three canonical metrics-SE, CD and LE. Spike times were converted into ISIs since our metrics probe the temporal relationship between successive spikes.

#### Task-relevant ISIs

For each neuron, we extracted ISIs during the *VS* and *MS* phases by filtering trials according to the `stim phase` key. Within each selected trial, spike times were clipped to the interval [`start time, stop time`]. To avoid artificial ISIs spanning trial boundaries, we computed intervals separately for each trial and then concatenated them across all trials of the same phase. Neurons with fewer than 20 ISIs in a given phase were excluded, as reliable estimation of chaos metrics requires a minimal sequence length for stable embedding, robust entropy estimation, and valid surrogate comparisons [13, 14] employed in this study. This conservative threshold excludes under-sampled sequences while retaining sufficient data for population-level analysis. Finally, ISIs were normalized to remove firing-rate differences, standardize scale for embedding parameters, reduce bias in entropy and dimension estimates. Since affine transformations do not alter the presence or absence of chaotic dynamics, this step allows robust classification of recordings as chaotic versus non-chaotic without affecting qualitative outcomes.

#### Task-irrelevant ISIs

To quantify intrinsic neural dynamics in the absence of task engagement, we extracted task-irrelevant ISIs from temporal gaps between annotated trials. These inter-gap periods were identified by locating non-overlapping segments in the full session timeline. ISIs were calculated separately within each continuous inter-gap segment, ensuring that intervals were not artificially combined across discontinuous gaps. The same filtering rules applied: sequences with fewer than 20 ISIs were excluded and these filtered sequences were normalized as outlined above before metric computations.

### Chaos detection pipeline

To quantify dynamical complexity in the normalized ISI sequences from the *VS, MS*, and *TI* phases, we computed three complementary chaos metrics: the sample entropy (SE), correlation dimension (CD), and largest Lyapunov exponent (LE). These metrics capture distinct, theoretically grounded aspects of nonlinear dynamics:

- *SE* reflects the temporal unpredictability of ISI sequences by quantifying the likelihood that similar spike patterns remain similar in the future;
- *CD* estimates the fractal dimensionality of the attractor governing the system’s state evolution;
- *LE* measures exponential sensitivity to initial conditions, a hallmark of chaotic systems.

The procedures used to compute each of the above chaos metric are described below and the surrogate statistical testing framework is detailed in the subsequent section.

#### Sample entropy (SE)

We computed SE to quantify the temporal unpredictability of ISI sequences [14]. For embedding dimension *m*, we defined vectors:

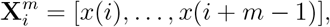

and used the Chebyshev norm to assess similarity:

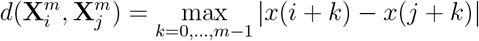

SE was defined as:

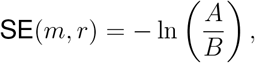

where *B* is the count of matching pairs of length *m* and *A* is the count of matching pairs of length *m* + 1, under a tolerance *r*.

To ensure robust estimation, we performed parameter sweeps over *m ∈* {2, 3} and *r ∈* {0.15, 0.2, 0.25} *×* SD(*x*). SE was computed for all (*m, r*) combinations, and the maximum SE value observed over this grid was reported as the final estimate for each ISI sequence.

#### Correlation dimension (CD)

We computed CD using a variant of the Grassberger–Procaccia algorithm [15], which estimates the dimensionality of the underlying attractor in reconstructed state space. For each embedding dimension, we constructed the matrix of pairwise distances between embedded vectors. Temporal correlations were suppressed by excluding pairs within a Theiler window of *T*_Theiler_ = 10 samples.

The correlation sum *C*(*r*) was computed as the proportion of temporally de-correlated embedded vector pairs separated by distance less than *r*:

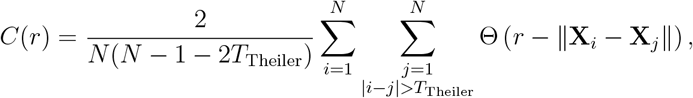

where Θ is the Heaviside function. We evaluated *C*(*r*) across 25 logarithmically spaced values of *r*, spanning the 5th to 95th percentiles of the distance distribution. CD was estimated as the local slope of log *C*(*r*) vs. log *r* over the central 50% of this radius range. To ensure robust estimation, CD was computed across all embedding dimensions *m ∈* {2, 3, 4, 5}, and the maximum slope observed over this grid was reported as the final CD estimate for each ISI sequence.

#### Largest Lyapunov exponent (LE)

We estimated the LE using the algorithm of Wolf et al. [16], which tracks the divergence of initially nearby trajectories in reconstructed state space. Delay embedding was performed as:

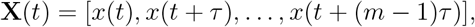

where *m* is the embedding dimension and *τ* is the delay parameter. For each reference vector **X**(*t*_*i*_), a temporally de-correlated nearest neighbor **X**(*t*_*j*_) was selected such that |*t*_*i*_*−t*_*j*_| *> T*_Theiler_ to avoid autocorrelated matches.

At each time step, we tracked the Euclidean distance between the evolving pair of trajectories. Let 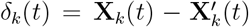denote the vector difference between the reference and its nearest neighbor at step *t*. The local divergence was evaluated over a single step Δ*t* = 1, and the largest Lyapunov exponent was estimated as:

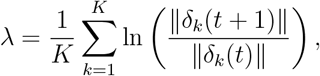

where *K* is the number of valid trajectory pairs tracked.

To ensure robust estimation, we performed parameter sweeps over *m ∈* {3, 4, 5, 6} and *τ ∈* {1, 2, 3}. For each ISI sequence, LE was computed across all combinations of (*m, τ*), and the maximum value observed was retained as the final estimate for that neuron.

The Wolf algorithm yields only non-negative values: values near zero indicate periodic or quasi-periodic dynamics, intermediate positive values suggest chaotic behavior, and larger values are typically observed in stochastic or noise-dominated signals.

### Surrogate Generation

In order to distinguish structured nonlinear or chaotic dynamics from stochastic variability i.e., noise, phase-randomized surrogate testing [17] was employed. In general, each metric depends sensitively on algorithmic parameters (e.g. embedding dimension, tolerance, reconstruction delay), intrinsic timescales, sampling rate and attractor geometry. Consequently, different signals yield a range of metric magnitudes even if they have same underlying temporal structure making classification based solely on absolute metric values infeasible. Thus, surrogate testing provides a statistically robust classification of chaos versus noise. Moreover, combining multiple surrogate-normalized features into a multivariate classifier enhances sensitivity and specificity. Parameter-sweep analyses (e.g. varying embedding dimensions and tolerances) further ensures that detected signatures of chaos persist across reconstruction settings rather than arising from narrow, potentially artifactual regimes.

For each ISI sequence, 100 surrogate time series were generated by preserving the

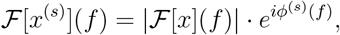

where the phases *ϕ*^(*s*)^(*f*) were drawn uniformly from [0, 2*π*] and adjusted to ensure Hermitian (conjugate) symmetry, thereby ensuring real-valued inverse transforms. These surrogates preserve all linear statistics but destroy higher-order dependencies, serving as a baseline: genuine chaotic dynamics will produce lower entropy, correlation dimension and divergence in the raw data than in its surrogates.

Each chaos metric was then recomputed on the individual surrogate sequences using the same parameter sweep with the maximum value recorded per metric in addition to the raw sequence as specified above. Any ISI sequence that resulted in undefined, infinite, or NaN values during metric computation was excluded to ensure validity and interpretability of downstream results. Statistical classification using surrogates is detailed later in this section.

### Testing periodicity

While the phase-randomized surrogate testing method confirms that neural activity is not stochastic, possibility remains that temporal structure could be chaotic or periodic. To rule out possible periodicity in the neuronal ISI sequences, we performed an autocorrelation function (ACF) analysis across all the neuronal data. The ACF quantifies temporal dependencies by measuring similarity between the raw ISI sequence and lagged copies of itself:

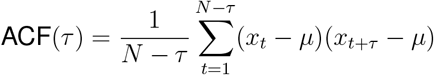

where *τ* denotes the lag, *µ* is the mean of the sequence, and *N* is the total number of ISIs. Each ACF was normalized such that ACF(0) = 1. The maximum lag was set to five seconds to ensure detection of slow rhythmic patterns if present.

The ACF computed for each ISI sequence was averaged across all recorded neurons, and this population-averaged ACF showed a strong peak at zero lag followed by a rapid decline to near-zero values with no evidence of evenly spaced secondary peaks throughout the entire lag range examined. In a periodic signal, the ACF would exhibit regularly spaced peaks at fixed lag intervals that reflect its rhythmic structure. The absence of such a pattern in the present data suggests that the neural activity does not exhibit periodicity.

### Statistics: Chaos Analysis

This subsection describes the statistical and comparative analyses performed after computing chaos metrics. These include criteria for significance testing, distributional comparisons, effect size estimation, and qualitative visualization techniques that support the main results.

#### Significance testing

For all the metrics (SE, CD, and LE), we assessed how often surrogate data exhibited greater apparent irregularity, complexity or divergence compared to the raw or original data by applying a one-sided test in a uniform direction across metrics [18].

Considering that some cases of surrogate distributions could be non-Gaussian, we avoided parametric assumptions and based all significance testing exclusively on empirical *p*-values.

The `empirical` *p*-value was computed as:

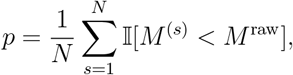

where 𝕀 [*·*] is the indicator function, equal to 1 if the condition is true and 0 otherwise, *M*^(*s*)^ denotes the metric value for the *s*-th surrogate, and *M* ^raw^ is the metric value computed from normalized ISI sequence prior to surrogate generation (referred to as raw metric value in this article). A metric computation was considered significant if `empirical` *p <* 0.025, corresponding to the raw metric value falling in the lowest 2.5% of the surrogate distribution. This left-tailed test reflects our expectation that *lower* values of SE, CD, and LE indicate more regular, lower-dimensional, and predictable dynamics than those generated by random fluctuations.

Note that the chaos metrics chosen for this study were based on their theoretical complementarity and their relevance to characterizing deterministic chaos in time series. While each provides a different perspective namely divergence, geometric complexity, or temporal irregularity, none is sufficient on its own to reliably distinguish chaos from stochastic variability. We therefore adopted a conservative criterion: only neurons showing significant deviations from surrogate distributions on all three metrics were classified as chaotic.

#### Benchmarking

To validate our analysis pipeline we applied the full procedure including surrogate testing, to synthetic signals with known dynamical properties. This ensured that our framework could detect nonlinear temporal dynamics when present thereby differentiating between chaotic and stochastic structures.

As a control signal, we generated 1000 samples of white noise by drawing independently from a standard normal distribution. This signal lacks temporal structure and served as a negative control. For the positive control, we simulated the logistic map, a known chaotic system, using the recurrence relation

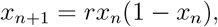

with *r* = 3.99 and initial condition *x*_0_ = 0.5, iterated for 1000 steps. These values produce a chaotic trajectory densely filling the unit interval.

For completeness, we ran the metrics on a synthetic periodic signal. For that purpose, we generated a smooth periodic signal by evaluating sin(*t*) over 1000 evenly spaced points span-ning the interval *t ∈* [0, 4*π*]. Although temporally structured, this sine wave contains no chaotic dynamics.

Each of these synthetic time series was z-scored. The resulting sequence was treated as a pseudo-ISI series and processed identically to the neuronal data. We computed the three chaos metrics for each series. For SE, we swept over embedding dimensions *m ∈* {2, 3} and radius parameters *r ∈* {0.15, 0.2, 0.25}·std(*x*) and retained the maximum value. CD was estimated using embeddings with *m ∈* {2, 3, 4, 5}. LE was estimated by sweeping over embedding dimensions *m ∈* {3, 4, 5, 6} and delays *τ ∈* {1, 2, 3}.

To assess significance, we generated 100 phase-randomized surrogate sequences for each synthetic signal. Metrics were recomputed on each surrogate, and the raw metric value was compared against the surrogate distribution using one-tailed empirical *p*-values.

#### Demographic effects

We tested the effect of gender on the overall percentage of chaotic neurons in each participant using Mann–Whitney U test, a non-parametric test appropriate for the observed non-normality of metric distributions. Additionally, Spearman correlation coefficient was estimated to understand the relationship between age and percentage of chaotic neurons.

#### Strength of chaos

To characterize the consistency and variability of nonlinear modulation across individual neurons, we computed the difference between each neuron’s raw metric value and its corresponding surrogate mean (Δ = raw *− µ*_surr_) for SE, CD, and LE. The resulting Δ distributions provide a neuron-by-neuron quantification of deviation strength from surrogate-based expectations, complementing the population-level shifts reported in Mann–Whitney U test. For each metric, we plotted the full distribution of Δ values separately for task-relevant and task-irrelevant phases.

To quantify the strength of nonlinear structure within passing neurons, we also computed z-scores for each chaos metric, defined as the deviation of the raw metric value from the surrogate mean, normalized by the surrogate standard deviation. These z-scores were computed from surrogate distributions that were not assumed to be Gaussian; rather, they served purely as standardized measures of the strength and magnitude of deviation relative to surrogate expectations, without being used for significance testing.

#### Phase-wise differences

Differences in chaos metrics between task-relevant (*VS* and *MS*) and task-irrelevant conditions were evaluated using two-sided Mann–Whitney U tests. For comparison of chaotic dynamics between any two task phases, we computed effect sizes using Cliff’s delta (*δ*), a non-parametric measure robust to skewed or non-normal distributions [19] and CLES, which quantifies the probability that a randomly selected value from one group exceeds that from another [20]. Cliff’s delta, with interpretive thresholds were derived from Vargha & Delaney [21] and its mapping to *δ* via *δ* = 2*A−*1. We approximate the small, medium, and large effect size boundaries to |*δ*| > 0.15,> 0.33, > 0.47, respectively, in line with common convention and practice in the field. In addition, CLES of 50% was interpreted as complete overlap (i.e., no effect), while values approaching 100% (or 0%, depending on directionality) reflect stronger effects. These statistics provided an interpretable, distribution-free quantification of changes in nonlinear dynamics across behavioral states.

Effects of demographics on the chaotic dynamics between task-relevant and task-irrelevant cases were investigated using the same methods outlined earlier (see Demographic effects).

#### Regional effects

Final step of our study involved probing any differences in the signatures of chaos across recorded neurons in the context of various brain regions. The dataset includes recordings from the amygdala and hippocampus from both left and right brain hemispheres.

To evaluate any differences in the counts of chaotic neurons between these four brain areas, we used the Friedman test. Moreover, influence of demographics on chaotic activity across these brain areas was quantified using methods outlined earlier (see Demographic effects).

## Results

Chaotic dynamics may shape memory function by decoupling current activity from long-past trajectories, thereby introducing flexibility into an otherwise presumably stable network. Since we analyze data recorded from brain areas implicated with a role in memory, we proposed that some fraction of MTL neurons would exhibit signatures of chaos. To test this, we developed and applied a systematic ISI-based protocol for distinguishing deterministic chaos from purely stochastic variability as outlined in *Methods*. We present the corresponding results in this section.

### Benchmarking

Before applying our chaos detection pipeline to neuronal recordings, we benchmarked metric outputs using raw or original and surrogate synthetic signals that are known to be stochastic, chaotic and periodic. For a high-dimensional *white* noise, we obtained SE = 2.75, CD = 2.90 and LE = 1.72, with no statistically significant deviation between raw and phase-randomized surrogate values (`empirical` *p >* 0.6). This confirms that our metrics yield high values for stochastic signal and that surrogate testing itself does not introduce artifactual divergence.

In contrast, purely chaotic signals produced lower raw metric values and clear divergence from their surrogate counterparts. For instance, using the logistic map we obtained raw metric value for SE = 0.61 versus surrogate mean (*µ*_surr_) = 2.54, CD = 1.95 versus *µ*_surr_ = 2.80, and LE = 0.94 versus *µ*_surr_ = 1.72 (`empirical` *p <* 0.025 for each metric). Thus, phase-randomized surrogates break any deterministic temporal structure, rendering them statistically indistinguishable from white noise.

While we ruled out the presence of periodicity in the neuronal data, we applied these three metrics to known periodic signals for completeness. These computations lead to raw metric values much lower than those for the synthetic chaotic data reported above. For instance, using a sine wave as a signal of interest, computing the raw metrics leads to values of 0.036 for SE,0.85 for CD and 0.038 for LE.

Overall, benchmarking and surrogate testing suggests that our metrics are capable of distinguishing between chaotic temporal structure versus unstructured noisy signals. For instance, applying the metrics to phase-randomized surrogates of chaotic signal results in mean values similar to those obtained from stochastic data. Moreover, the raw metric values from this chaotic signal are significantly smaller than those from corresponding surrogates suggesting presence of deterministic chaos.

### Chaotic Neurons

Applying all three metrics to our filtered set of task-relevant neurons revealed that a substantial proportion deviated significantly from their phase-randomized surrogates, consistent with signatures of chaotic dynamics. Out of the 1863 neurons in the dataset, after excluding short ISI sequences and failed metric computations, during the *VS* phase, 91.97% (1180 of 1283) passed significance (`empirical` *p* < 0.025) for SE, 90.60% (1244 of 1373) for CD, and 58.06% (886 of 1526) for LE. In the *MS* phase, 98.15% (1330 of 1355) passed for SE, 98.20% (1330 of 1355) for CD, and 57.45% (936 of 1629) for LE. Furthermore, during the *TI* period, many neurons still showed significant deviations: 55.53% (306 of 551) for SE, 99.83% (603 of 604) for CD, and 66.67% (458 of 687) for LE. These results demonstrate that chaotic temporal structure is present across behavioral states, with stronger and more consistent signatures emerging during active memory engagement.

However, to conservatively classify a neuron as chaotic, we required it to exhibit statistical significance across all three metrics simultaneously. Quantitatively, we successfully computed the metric values for 1242 overlapping neurons between all three metrics out of which 790 or 63.60% passed significance in *VS* phase. Similarly, 827 out of 1342 or 61.62% neurons passed significance on all the three metrics simultaneously in the *MS* phase. In contrast, during the *TI* phase, 263 out of 551 or 47.73% passed all three metrics, with neuronal overlaps correspondingly reduced. This pattern suggests that coherent nonlinear structure preferentially emerges as a multi-metric phenomenon during behaviorally engaged phases. Figure 2A-C visualizes this overlap of neurons with individual metrics passing or failing statistical significance, while Figure 2D shows distribution of the raw metric values for neurons classified as chaotic and non-chaotic.

**Figure 2:**
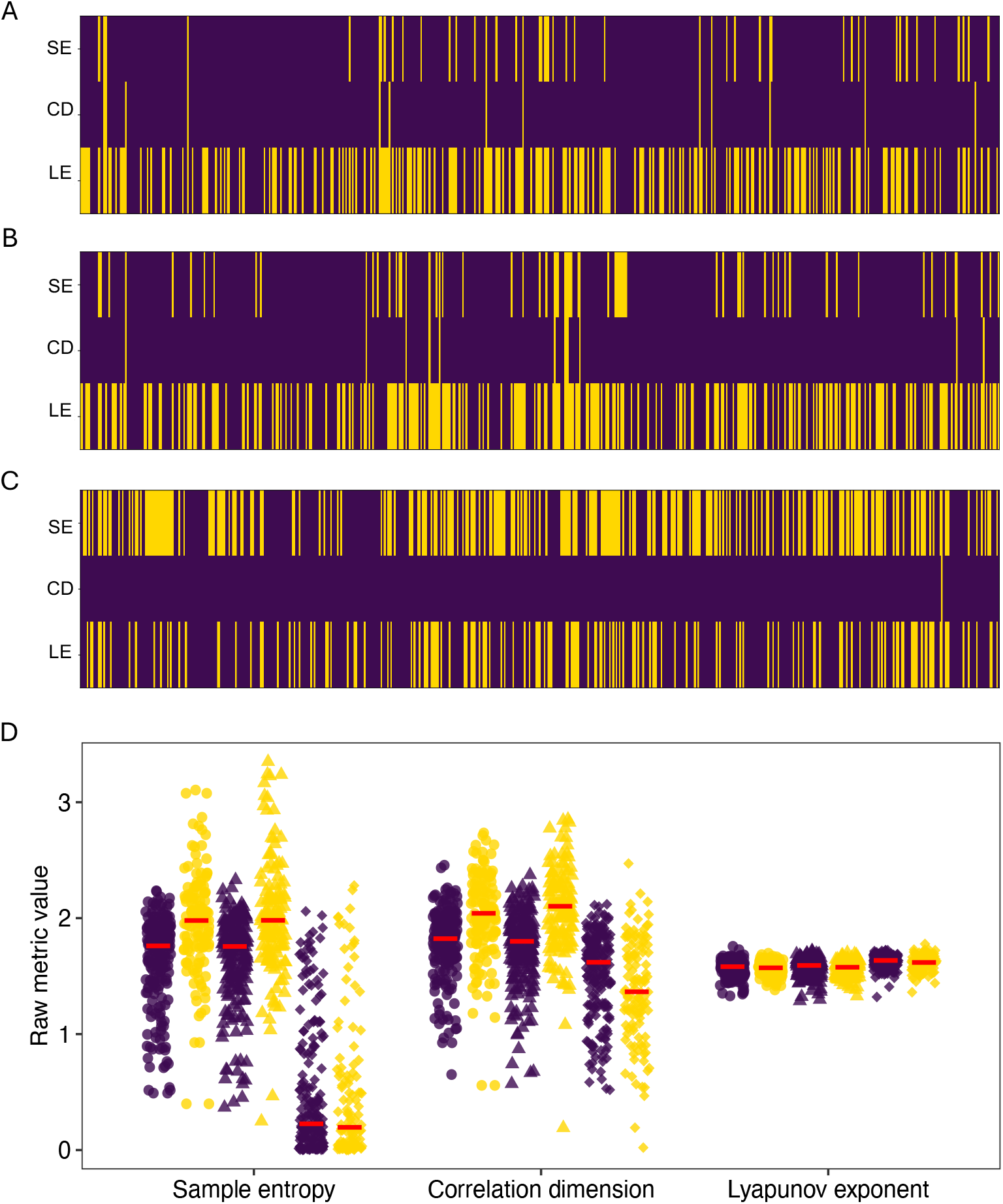
Visualizing distribution for chaotic and non-chaotic neurons during A: visually-selective, B: memory-selective, task-irrelevant phases, and D: based on raw metric values. Dark purple and yellow markers indicate statistical significance and insignificance, respectively. In A, B and C continuous vertical dark purple line over the extent of each plot indicates neurons that passed all three metrics simultaneously. In D, Circles, triangles and diamond markers represent visually-selective, memory-selective and task-irrelevant phases, respectively. Red bars indicate median values of the distributions.

Additional assessment at the participant level involved computing the proportion of neurons exhibiting evidence of chaos out of the total number of neurons recorded from that individual. This analysis revealed no significant difference in the overall percentage of chaotic neurons between males and females (Mann-Whitney U = 361.5, *p* = 0.30) and there were no correlations found between age and percentage of chaotic neurons (`Spearman’s` *ρ* = 0.10, *p* = 0.46).

Thus far, we have classified neurons as chaotic based on whether their raw metric values significantly deviated from the distribution of values obtained from 100 phase-randomized surrogates. Now, in order to probe the extent of this deviation, we compute the difference (Δ) between raw metric value and surrogate mean per neuron for each metric and plot the distributions for the *VS, MS* and *TI* phases (Figure 3). The majority of Δ values are negative, indicating that raw metric values fall below respective surrogate means, consistent with deterministic temporal structure in the neuronal activity.

**Figure 3:**
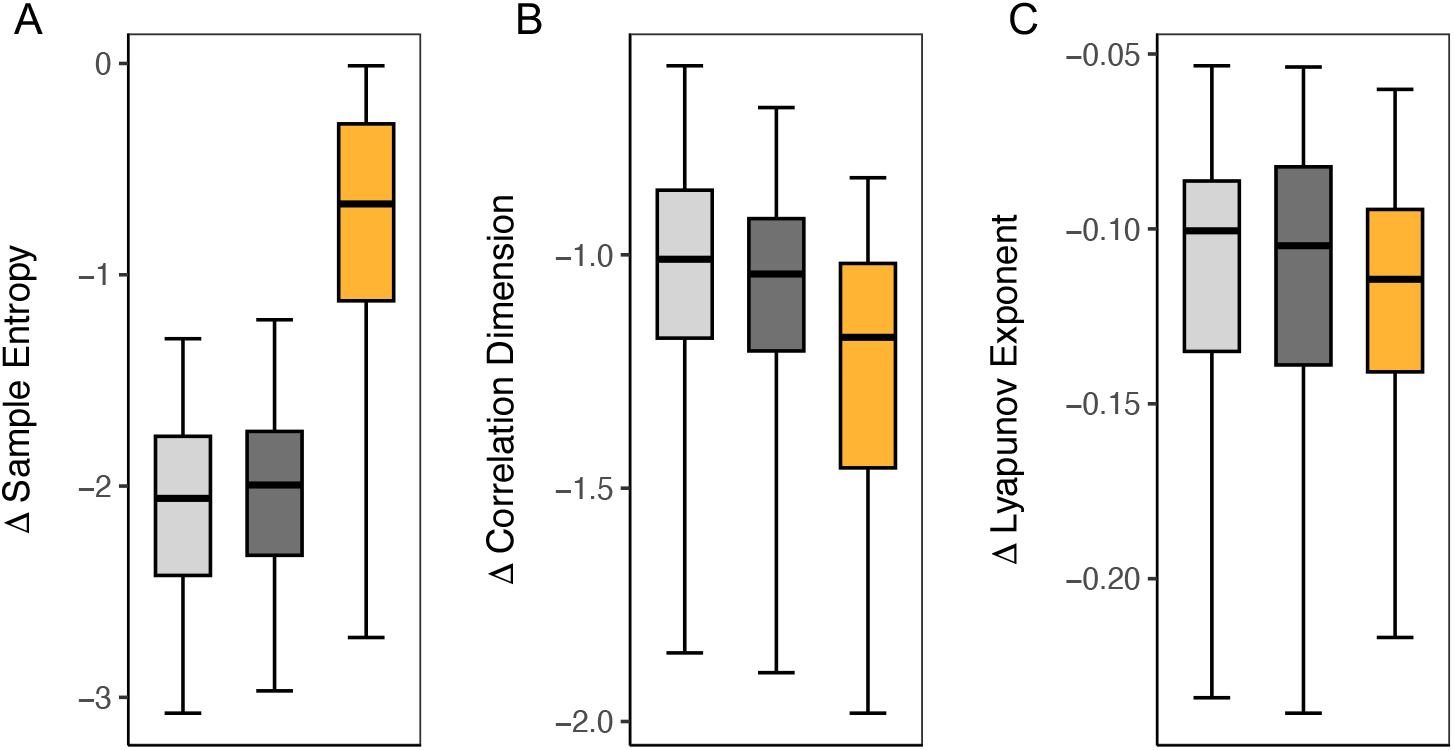
Boxplots of Δ = raw *µ*_surrogate_ for A: sample entropy, B: correlation dimension, and C: the largest Lyapunov exponent across visually-selective (light gray), memory-selective (dark gray) and task-irrelevant (orange) phases. Negative Δ indicates metric values below surrogate means, suggesting deterministic or chaotic structure.

Furthermore, we computed z-scores of these differences for neurons passing significance on all three metrics to quantify the average *strength* of chaos. Across neurons that passed significance, the population z-scores are *−*6.66 for SE, *−*20.34 for CD and *−*5.12 for LE, indicating highly significant deviations (|*z*| > 3). Thus, the deviation between raw metric values and surrogate means is statistically significant and not a result of jitter or any other artifactual fluctuations.

### Task-Relevant versus Task-Irrelevant

The metric computations on raw and surrogate ISI sequences yield a wide range of values. Table 1 summarizes the mean and median raw metric values of SE, CD and LE for both task-relevant (*VS* and *MS*) and irrelevant conditions, alongside values from corresponding surrogates.

**Table 1:** Mean and median raw metric values for visually-selective, memory-selective, and task-irrelevant phases for all three metrics significant (`empirical` *p* < 0.025). SE: sample entropy, CD: correlation dimension, and LE: largest Lyapunov exponent.

| Metric | Visually-Selective |  | Memory-Selective |  | Task-Irrelevant |  |
| --- | --- | --- | --- | --- | --- | --- |
|  | Raw | Surrogate | Raw | Surrogate | Raw | Surrogate |
| SE (Mean) | 1.57 | 3.60 | 1.63 | 3.53 | 0.48 | 1.20 |
| SE (Median) | 1.65 | 3.56 | 1.70 | 3.50 | 0.20 | 0.92 |
| CD (Mean) | 1.72 | 2.83 | 1.74 | 2.83 | 1.52 | 2.82 |
| CD (Median) | 1.76 | 2.83 | 1.79 | 2.83 | 1.61 | 2.84 |
| LE (Mean) | 1.58 | 1.69 | 1.59 | 1.70 | 1.63 | 1.75 |
| LE (Median) | 1.59 | 1.70 | 1.60 | 1.71 | 1.64 | 1.76 |

Interestingly, SE surrogate values exceed those of white noise (see Benchmarking) in task-relevant phases (*VS* and *MS*). This perhaps arises because task-relevant ISIs often contain rhythmic or burst-like patterns, resulting in structured power spectra with elevated low-frequency power. When phases are randomized in such signals, the resulting surrogates retain this smooth spectral shape but lack coherent structure in time thereby inflating the number of near-matches used in entropy calculation and pushing SE above that seen in white noise.

In contrast, surrogate values for SE during *TI* phase are lower than that for white noise. It is possible that the corresponding ISIs often contain uneven spike timing that yields jagged or less structured power spectra. Phase randomization of such signals does not produce temporally smooth or entropy-rich sequences, leading to surrogate SE values even lower than that for white noise. This makes SE particularly sensitive to biologically relevant dynamics.

Nevertheless, the variability in surrogate SE across phases does not compromise the validity of the significance testing framework as is also evident from the computations using synthetic chaotic signal (discussed earlier in Benchmarking. Regardless of the surrogate’s absolute SE value, the key comparison is always relative: whether the raw SE is significantly lower than its own surrogates. This ensures that each ISI sequence is evaluated against an appropriate baseline that preserves its spectral properties.

On the other hand, CD and LE, both geometry-based measures, yield similar metric values for surrogates and white noise, since both destroy the attractor’s true deterministic geometry. Overall, we see that raw metric values for the task-relevant phases are consistently higher (for SE and CD) or lower (for LE) than those for task-irrelevant phase. To quantify these differences, we conduct a task phase-wise analysis as described in the following section.

### Phase-wise differences

Behavior-dependent shifts in chaos metrics were assessed by comparing the three task phases (*VS, MS* and *TI*) using two-sided Mann–Whitney *U* tests. To ensure a fair comparison, we restricted our analyses to neurons that pass significance on all three metrics and simultaneously occur in the phases being contrasted: 171 neurons for *VS* vs. *TI*, 164 neurons for *MS* vs. *TI*, and 604 unique neurons between the *VS* and *MS* phases. Notably, analyses conducted on the full set of phase-specific data yielded qualitatively similar results.

All three chaos metrics differed with high significance in both *VS* vs. *TI* and *MS* vs. *TI* comparisons (*p* < 0.001). SE was elevated during task-relevant phases, reflecting increased temporal irregularity; CD was higher, indicating a more complex attractor structure; and LE was reduced, consistent with more constrained, predictable dynamics under cognitive engagement.

To quantify effect sizes, we computed Cliff’s delta (*δ*) and the common-language effect size (CLES) for each contrast (see Table 2). Cliff’s delta (*−*1 *≤ δ ≤* 1) is interpreted as small for |*δ*| > 0.15, moderate for |*δ*| > 0.33, and large for |*δ*| > 0.47. CLES represents the probability that a randomly selected value from one subset exceeds (for SE and CD) or is lower (for LE) than a value from another.

**Table 2:** Effect sizes for chaos metrics across visually-selective (VS), memory-selective (MS) and task-irrelevant phases. Cliff’s *δ* quantifies distributional non-overlap; Common-language effect size (CLES) gives the probability that a value from the first condition exceeds (for sample entropy and correlation dimension) or is lower (for largest Lyapunov exponent) than a value from the second condition. p-values are from two-sided Mann-Whitney U tests.

| Comparison | Metric | Cliff's $\delta$ | CLES % | p-value (Mann-Whitney) |
| --- | --- | --- | --- | --- |
| *VS vs TI* | SE | 0.87 | 93.87 | < 0.001 |
|  | CD | 0.38 | 69.02 | < 0.001 |
|  | LE | -0.54 | 23.08 | < 0.001 |
| *MS vs TI* | SE | 0.87 | 93.98 | < 0.001 |
|  | CD | 0.29 | 64.87 | < 0.001 |
|  | LE | -0.45 | 27.65 | < 0.001 |
| *VS vs MS* | SE | -0.04 | 47.97 | 0.22 |
|  | CD | -0.026 | 48.68 | 0.42 |
|  | LE | -0.12 | 44.02 | < 0.001 |

SE showed large positive modulation during task engagement (*δ >* 0.8, CLES > 93%), indicating that in over nine out of ten random neuron pairs, the task-relevant neuron has higher SE than the task-irrelevant one. On the other hand, CD exhibited small to moderate effects (*δ ∼*0.3, CLES *∼*65%), suggesting elevated attractor dimensionality during task-relevant vs. irrelevant conditions. The CLES indicates that over six out of ten random neuron pairs will have CD higher for task-relevant vs. irrelevant scenarios. Lastly, LE showed moderate negative effects (*δ* between –0.45 and –0.54, CLES *∼*30%), implying that roughly two thirds of task-relevant neurons have lower LE and thus more chaotic (less noise like) dynamics than task-irrelevant neurons. The direct comparison between *VS* and *MS* phases revealed negligible differences for all the three metrics (|*δ*| *<* 0.15, CLES *∼*44–48%), although Mann-Whitney test revealed significant difference in LE for this contrast.

Overall, these results indicate that task engagement not only increases variability but also induces richer and more structured temporal dynamics, shifting neural activity into a chaotic regime with higher temporal complexity, greater dimensionality and lower divergence during task engagement. Moreover, this analysis was extended to account for any demographic effects namely participant age and gender.

While there were no significant differences in the percentage of chaotic neurons between males and females in the *VS* and *MS* phases, during the *TI* phase alone, males exhibited a higher percentage of chaotic neurons than females (U = 290.0 *p* = 0.017, Figure 4A-C). This suggests greater neural variability or reduced network stability in males when the task is not behaviorally relevant. Additionally, no significant age association was found with the percentage of chaotic neurons in the *VS* phase (*ρ* = 0.04, *p* = 0.76), *MS* phase (*ρ* = 0.10, *p* = 0.47) or the *TI* phases (*ρ* = 0.11, *p* = 0.42) (Figure 4D-F).

**Figure 4:**
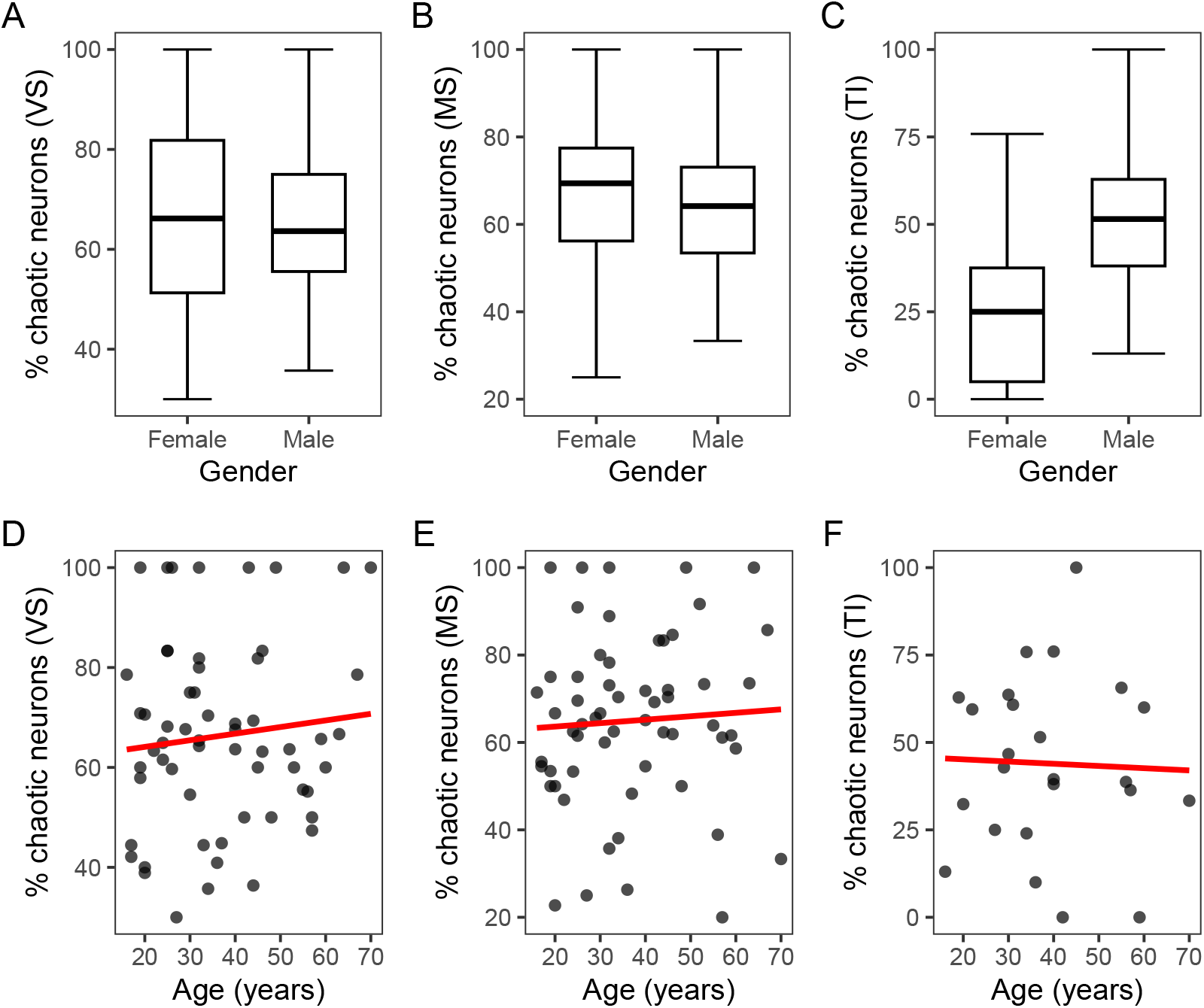
Percentage of triple chaotic neurons across gender in A: visually-selective, B: memory-selective and C: task-irrelevant phases. Association between age and percentage of chaotic neurons in D: visually-selective, E: memory-selective and F: task-irrelevant phases.

### Regional Effects

Finally, we examined whether chaotic dynamics varied by anatomical region. Neurons passing significance on all three metrics were grouped by hemisphere and structure-left amygdala, left hippocampus, right amygdala and right hippocampus. Figure 5A shows the counts of chaotic neurons in each of these regions for *VS, MS* and *TI* phases. Although the right hippocampus contributed the largest number of chaotic neurons, all four regions displayed substantial nonlinear structure. The Friedman test confirmed that regional differences with respect to number of chaotic neurons were not significant (*p >* 0.1), indicating that chaos is a widespread feature of MTL circuitry.

**Figure 5:**
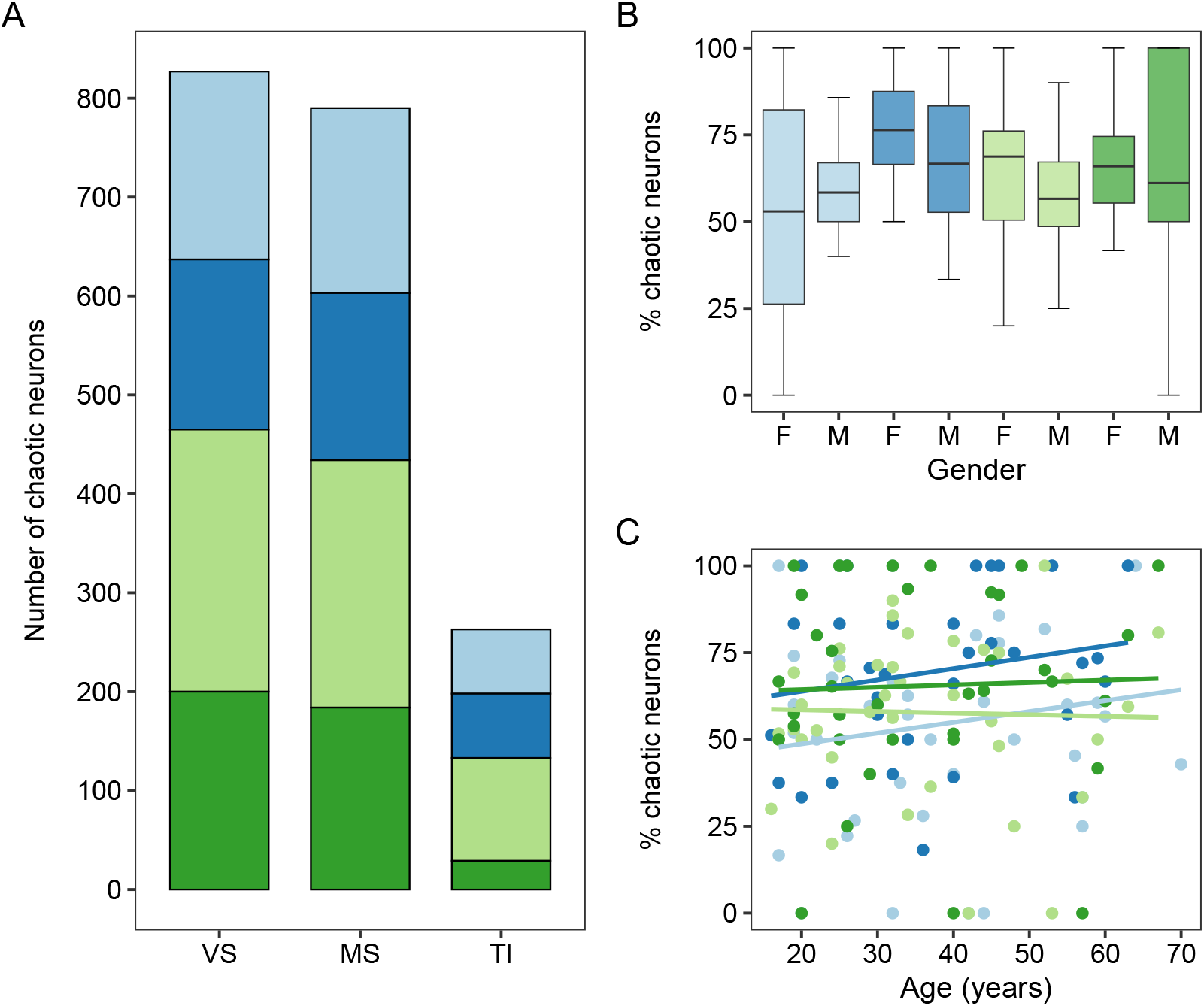
Regional breakdown of triple-significant neurons during visually-selective, memory-selective and task-irrelevant phases. A: Bars denote counts in left amygdala (light-blue), left hippocampus (dark-blue), right amygdala (light-green) and right hippocampus (dark-green), B: percentage of chaotic neurons across these regions with respect to gender, and C: percentage of neurons across the same regions with respect to age.

Region-specific analyses likewise showed no gender differences in left amygdala (U = 365.5, *p* = 0.32), left hippocampus (U = 362.5, *p* = 0.29), right amygdala (U = 381.0, *p* = 0.46) or the right hippocampus (U = 411.0, *p* = 0.78), nor age correlations in any region (all |*ρ*| *<* 0.24,*p >* 0.05) as shown in Figure 5B-C.

## Discussion

This study is a unique effort demonstrating robust signatures of chaotic dynamics in single neurons of the human MTL. Using surrogate-validated metrics we uncovered nonlinear or chaotic temporal structure that deviated from linear stochastic models. Crucially, this structure emerged across all three metrics: nearly 60% neurons in task-relevant and 48% neurons in task-irrelevant cases passed significance thresholds (`empirical` *p* < 0.025) on SE, CD and LE simultaneously, providing strong evidence for deterministic structure in spike timing.

Each metric contributed complementary insights. Elevated SE and CD captured increased temporal irregularity and attractor dimensionality, while reduced LE reflected tighter local constraints and greater predictability which is characteristic of chaotic regime. Requiring convergence across all three measures served as a conservative criterion for identifying low-dimensional chaos and helped rule out artifacts from individual metric biases or noise.

We use the term *low-dimensional chaos* based on theoretical and empirical grounds. Classical definitions describe such chaos as arising from deterministic systems with few interacting degrees of freedom, yielding structured but unpredictable dynamics [22, 13]. In our data, task-relevant neurons showed bounded SE, CD and LE, markedly lower than white noise and phase-randomized surrogates. This profile indicates that MTL dynamics lie on compact, interpretable manifolds rather than in high-dimensional or purely stochastic regimes.

Notably, chaotic signatures were modulated by behavior. During both *VS* and *MS* phases, SE and CD increased while LE decreased relative to *TI* periods, with significant effect sizes. LE showed a subtle additional reduction during the *VS* phase, perhaps imposing a slightly stronger constraint on trajectory evolution than visual recognition contingent on memory.

While our results demonstrate the presence of chaotic dynamics in individual neurons of the human temporal lobe, a key limitation is that we do not yet understand how these dynamics contribute to memory processes. In the absence of a comprehensive theoretical framework for memory function, immediate implications of chaotic neuronal activity remain unclear. Nonetheless, our findings lay the groundwork for future interdisciplinary investigations that may ultimately yield a unified, mechanistic account of whether chaotic dynamics actively support memory formation and retrieval or emerge as a byproduct of those processes. Moreover, such future efforts may also provide insights about whether single neurons generate chaotic dynamics intrinsically or these dynamics emerge from network inputs. Disentangling these possibilities may require intracellular recordings, network perturbations and modeling to determine the origin of low-dimensional chaos in neural circuits.

Another important consideration is that, although the data we analyzed come from epilepsy patients, the widespread anatomical distribution of chaotic signatures and their task-dependent modulation argue against a purely pathological origin. While seizure activity involves runaway firing rates, and could in principle alter chaos metrics, we presume that task-relevant data was recorded during interictal periods and excluded epochs with overt clinical seizures. However, we cannot fully rule out subclinical events during *TI* phases, yet the fact that chaotic dynamics are markedly less prevalent during these periods than during task engagement further undermines a direct link to epilepsy.

Together, these findings provide the first large-scale evidence that chaotic dynamics are an intrinsic feature of human single-neuron activity in the MTL and are selectively amplified during memory-guided behavior. Future work should explore how these dynamics are shaped by learning, cognitive load and neuromodulatory states, and whether they confer computational advantages such as generalization or memory indexing. Understanding chaos in single-neuron activity may also have translational value, as disruption of these dynamics could underlie deficits in memory disorders and targeted restoration might offer novel therapeutic avenues.

## Software and Reproducibility

All analyses were conducted in Python using standard scientific libraries, including NumPy, SciPy, pandas, and matplotlib. R version 4.5.0 was used for statistics and graphical representations. Phase-randomized surrogates were generated using an FFT-based algorithm that preserves the amplitude spectrum while randomizing phase components. ISI sequence extraction and full metric computation scripts are available at: https://github.com/neuro-research-codes/Chaos-Detection-Pipeline.git.

## Author Contributions

S.A. Chitale: Conceptualization, formal analysis, coding and computation, writing– original draft.

S. Ravichandran: Conceptualization, methodology, writing– review and editing.

## Competing Interests

The authors declare no competing interests.

## Acknowledgement

The authors gratefully acknowledge the resources provided by the University of Alabama at Birmingham IT-Research Computing group for high performance computing (HPC) support and CPU time on the Cheaha compute cluster. We also thank Chandravadia *et al*. for generating and curating the publicly available neuronal recordings from the declarative memory task.

## Notes

### Competing Interest Statement

The authors have declared no competing interest.

